# Phage metagenome-assembled genomes portraying anti-ESKAPE and anti-CRISPR/Cas potential; datasets from sewage-clinical settings of Western Uganda, sub–Saharan Africa

**DOI:** 10.1101/2025.03.12.642846

**Authors:** Jackim Nabona, Samweli Bahati, Abdalah Makaranga, Chinyere Nkemjika Anyanwu, R Neel, AbdulGaniy. B. Agbaje, Emmanuel Eilu, Deogratius Mark, Reuben S. Maghembe

## Abstract

ESKAPE pathogens include *Enterococcus faecium, Staphylococcus* aureus, *Klebsiella pneumoniae, Acinetobacter baumannii, Pseudomonas aeruginosa* and *Enterobacter* spp., which account for the major causes of mortality linked to the spread of infection and antimicrobial resistance (AMR) globally. Advances in omics approaches have pointed to bacteriophages as a promising alternative source of antibacterial agents. Here we enriched two samples from sewage and amplified them on *Staphylococcus* culture, followed by whole metagenome shotgun sequencing with Illumina NovaSe X. We performed metagenomic classification of high-quality sequence reads using the Kraken 2 database, to delineate the diversity and abundances of taxa. Thereafter we assembled the sequence reads with MAGAHIT and binned them with default parameters of the Bacterial and Viral Bioinformatics Resource Center (BV-BRC) before annotating each bin with PhageScope. From assembly, we recovered multiple metagenome-assembled genomes (MAGs) including *Alistipes* phage, *Escherichia* phage, *Vibrio* phage, *Staphylococcus* phage, *Klebsiella* phage and *Acinetobacter* phage, to mention the top six best hits. From annotation, while the *Acinetobacter* phage is virulent, the two *Klebsiella* phage and *Staphylococcus* phage are temperate. All the phages possess more than four lysis genes, with the potential to disrupt bacterial membranes. Exceptionally, *Vibrio* phage, *Acinetobacter* phage and *Alistipes* phage possess anti-CRISPR genes, the potential to counteract normal bacterial immune response to phage infection. These findings also inform that MAGs from the sewage have the potential to recover phages with anti-CRISPR/Cas activity, which is one of the desirable attributes for effective phage-bacterial infection to control the growth and multiplication of bacteria. Our datasets can be utilized for genome-guided selection of potent phages through lytic and host-range assays, towards the purification of endolysins (lysozymes) as alternative antibacterial agents.

**VALUE OF THE DATA:**

- Metagenome-assembled genomes (MAGs) of lytic phages could present a potential model to combat multidrug-resistant *Staphylococcus, Klebsiella and Acinetobacter* species, which are WHO’s high-priority pathogens.
- Phages with bacterial infective potential can be used as model gene vehicles and vectors for gene and genome editing studies as they possess hydrolytic enzymes targeting the bacterial cell walls, chromosomal DNA sequences and anti-CRISPR/Cas proteins.
- Comparing raw and processed datasets, MAGs provide an avenue for the pursuit of novel industrial strains from local resources in East Africa.

## BACKGROUND

ESKAPE includes *Enterococcus faecium, Staphylococcus* aureus, *Klebsiella pneumoniae, Acinetobacter baumannii, Pseudomonas aeruginosa* and *Enterobacter* spp, a group of the World Health Organization’s priority pathogens whose antimicrobial resistance (AMR) is a global threat. ESKAPE pathogens account for the largest AMR-associated mortality globally [1]. The evolution of AMR among these pathogens is drastic and has been experimentally shown to involve the selection of pre-existing bacterial variants [2] pre-empting the modes of action of preclinical antibiotic candidates of similar structural and functional relationships. This suggests an urgent need for novel antimicrobial scaffolds from different model sources. While phages have been under investigation for several decades, advances in omics technologies have rekindled the interest in the exploration of proteins with the lytic potential to control the growth and multiplication of bacteria including ESKAPE pathogens [3–5]. These datasets were generated on the assertion that ideal anti-ESKAPE phages should be isolated from similar environments to their hosts as they might have evolved to control the same bacteria. Thus, these metagenomic datasets were generated to prospect follow-up studies to establish lytic strains against multidrug-resistant members of the ESKAPE group.

## EXPERIMENTAL DESIGN, MATERIALS AND METHODS

### Sample collection, phage extraction, and enrichment

A total of 15 sewage water samples were aseptically collected from the top strata of flowing sewage water in sterilized Falcon tubes (50 ml) from the drainage systems of TASO and Kakoba Sewage Plants around River Rwizi (0°41′33″S 30°54′20″E). The samples were taken to the laboratory and stored at room temperature to settle any large and insoluble waste before taking the supernatant for phage isolation.

Samples from subclinical mastitis specimens retrieved from the Microbial Bank in the Department of Microbiology and Immunology at Kampala International University (KIU-WC) were used in bacteriophage isolation and propagation. The log phase *Staphylococcus* isolates were cultured in double strength (2X) tryptic soy broth (TSB) supplemented with 10mM MgSO4.7H2O and incubated at 37 ºC. 0.7% agar was prepared and used for making an overlay. Sewage samples were enriched using the method described by Beaudoin and colleagues [8] and modified according to Hyman [5] and with our minor modifications. We added 6 ml tryptone soy broth (TSB) and mixed it with an equal volume of sewage water and fresh *Staphylococcus* aliquot at log phase in a 250 ml conical flask. The phage-bacterial mixture was then incubated at 37°C in a mechanical shaker at 100 rpm overnight. Then the sewage-phage culture was centrifuged at 4,000 rpm for 10 minutes. The supernatant (bacteriophage-containing) was collected and filtered through a 0.45-micron filter. 10µl of the filtrate was spotted on to top agar layer of the TSA plate and allowed to dry before reintubation at 37 ºC overnight before isolation from plaques.

The obtained phage plaque stocks were amplified in TSB (broth culture) enriched with 10mM MgsO4 in their corresponding bacterial cell hosts that is *Staphylococcus* isolates grown overnight at (OD600 =04-0.6). These were incubated in a mechanical shaker at 70 rpm for 24 h at 37 ºC. Following lysis of the bacterial hosts, the bacteria-phage mixture was centrifuged at 5000 rpm for 5 min. The supernatant containing the phage was filtered using 0.45 µm syringe filters to get concentrated phage lysates that were transferred into new tubes before DNA extraction.

### DNA isolation, whole metagenome sequencing, and genome assembly

Metagenomic DNA samples were extracted from two of the samples using a Zymo Research miniprep kit as described in our previous work [9]. Libraries were constructed using a TruSeq Nano DNA (350) and sequenced using the Illumina NovaSeqX platform. All post-sequencing processes were performed in Python v3.10.14. Raw read sequences were then quality-controlled with FastQc (v0.12.1). The number of Fred scores of 20 and a maximum trimming error rate of 0.1. We then used default parameters of the Kraken 2 database within the Bacterial and Viral Bioinformatics Resource Center (BV-BRC, https://www.bv-brc.org/) to classify the metagenomic read sequences into the most appropriate taxa, whose abundances with computed with Bracken v1 [10]. Thereafter we employed the MEGAHIT (v1.2.9) package [11] to generate bins which are hereby defined as metagenome-assembled genomes (MAGs), which were then quality-assessed using machine learning with CheckM v2 [12].

### Metagenome-assembled genome annotation and phylogroups

Our interest was to recover more *Staphylococcus* phage-related genomes, we selected bins that were primarily matched with *Staphylococcus* phages. Then we selected the rest of the bins based on CheckM quality parameters and retrieved their comparative references from the NCBI Nucleotide Database for comparative phylogenetic analysis. The entire analysis was carried out by the VICTOR web service (https://victor.dsmz.de), a method for the genome-based phylogeny and classification of prokaryotic viruses [13]. All pairwise comparisons of the nucleotide sequences were conducted using the Genome-BLAST Distance Phylogeny (GBDP) method [14] under settings recommended for prokaryotic viruses [13]

The resulting intergenomic distances were used to infer a balanced minimum evolution tree with branch support via FASTME including SPR postprocessing [15] for each of the formulas D0, D4, and D6, respectively. Branch support was inferred from 100 pseudo-bootstrap replicates each. Trees were rooted at the midpoint [16] and visualized with ggtree [17]

Taxon boundaries at the species, genus, and family level were estimated with the OPTSIL program, the recommended clustering thresholds, and an F value (fraction of links required for cluster fusion) of 0.5 [13,14].

The bins with a completeness of above 98% and less than 5% contamination were annotated with PhageScope v1.0 [18] to predict taxonomic, host, virulence, lifestyle, and functional assignments from coding sequences (CDS). Other features such as genome size (bp), GC content (%), open reading frames (ORFs) mediating transcription termination, transmembrane protein expression, and anti-CRISPR activity were also recovered from the PhageScope annotation algorithms.

## DATA OVERVIEW

The datasets in this article describe the diversity and abundance of phage species characterized by a viral metagenome enriched by overnight cultivation on *Staphylococcus* sp. The Illumina NovaSeqX platform sequencing the first sample RSMJ3B3Sa generated a total of 18,246,958 raw reads with an average length of 151 bp each (equivalent to 1,377,645,329 bp). After quality control, the number of reads was reduced to 10,798,626 (1,464,449,844 bp). The second sample RSMJ51Sa yielded a total of 18,246,958 raw reads, equivalent to 1,377,645,329 bp, which upon filtration were reduced to 10,798,626 (1,371,106,335 bp). From Kraken 2 metagenome taxonomic analysis sample RSMJ3B3Sa two distinct genera of viruses namely *Baoshanvirus* and *Gamaleyavirus*.

Bracken Figure 1 (a) represents the largest number of classified sequences, which matched *Klebsiella* phage Miami (Virus), followed by *Staphylococcus cohnii* (Bacteria). Fig. 1(b) presents the abundances of each identified species in sample RSMJ51Sa, in which the highest sequence abundance was that of Achromobacter sp. AONIH (Bacteria) followed by *Gamaleyavirus* APEC7 (Virus) and then *Baoshanvirus* BS2 (Virus).

**Fig. 1.**
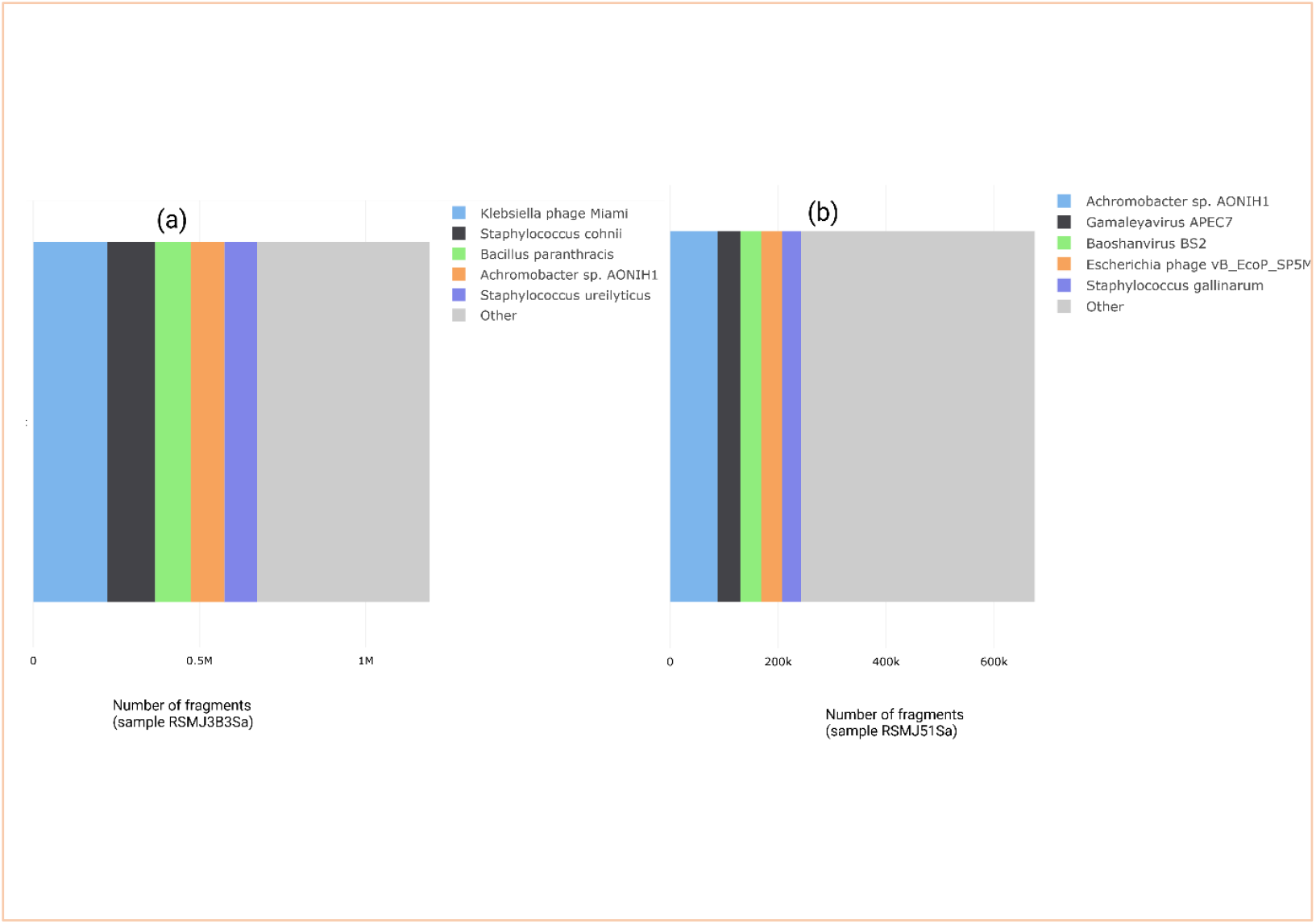
The relative abundances of the top five taxa at the species level were estimated using Bracken, based on high-quality read sequences. Below each bar plot, the number of DNA fragments accurately classified into their respective taxonomic groups is indicated.

Tables 1 and 2 represent MAGs with their preliminary matches and other features generated from BV-BRC-binning and CheckM assessment of the samples RSMJ3B3Sa and RSMJ51Sa, respectively. Table 1 shows that high-quality MAGs with above 95 % completeness and less than 5% contamination were bin 3, corresponding to *Staphylococcus* phage VB-SauS-SA2, (GenBank Accession GCA_003307435.1), bin 4, which matches with *Erwinia* phage vB_EamM_Joad (Accession GCA_002629065.1), bin 5, 2, and 7, which were not matched with known NCBI phage genome.

**Table 1.**
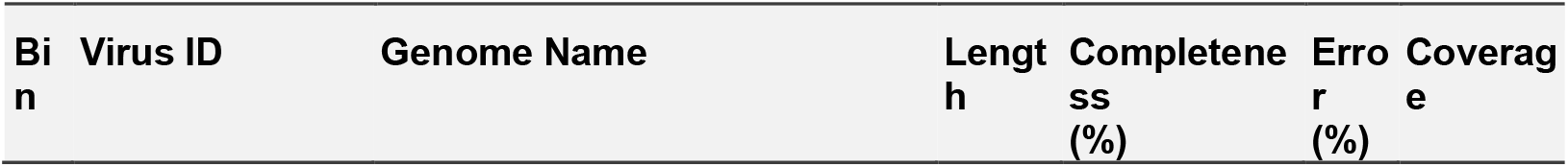
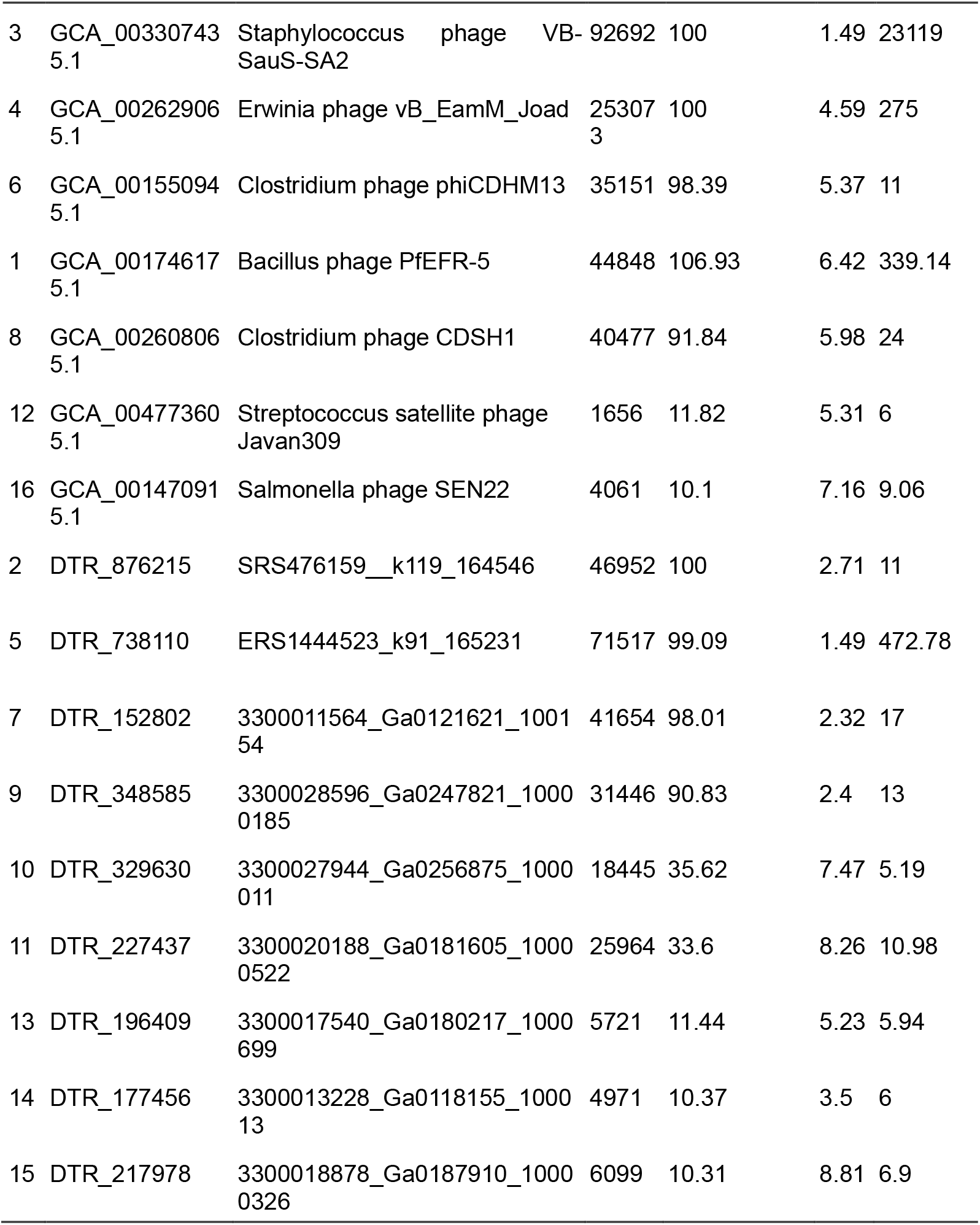
Binning results for sample RSMJ3B3Sa. Parameters including possible relative strain hit, bin size (length), completeness and other features are presented in corresponding columns.

**Table 2.**
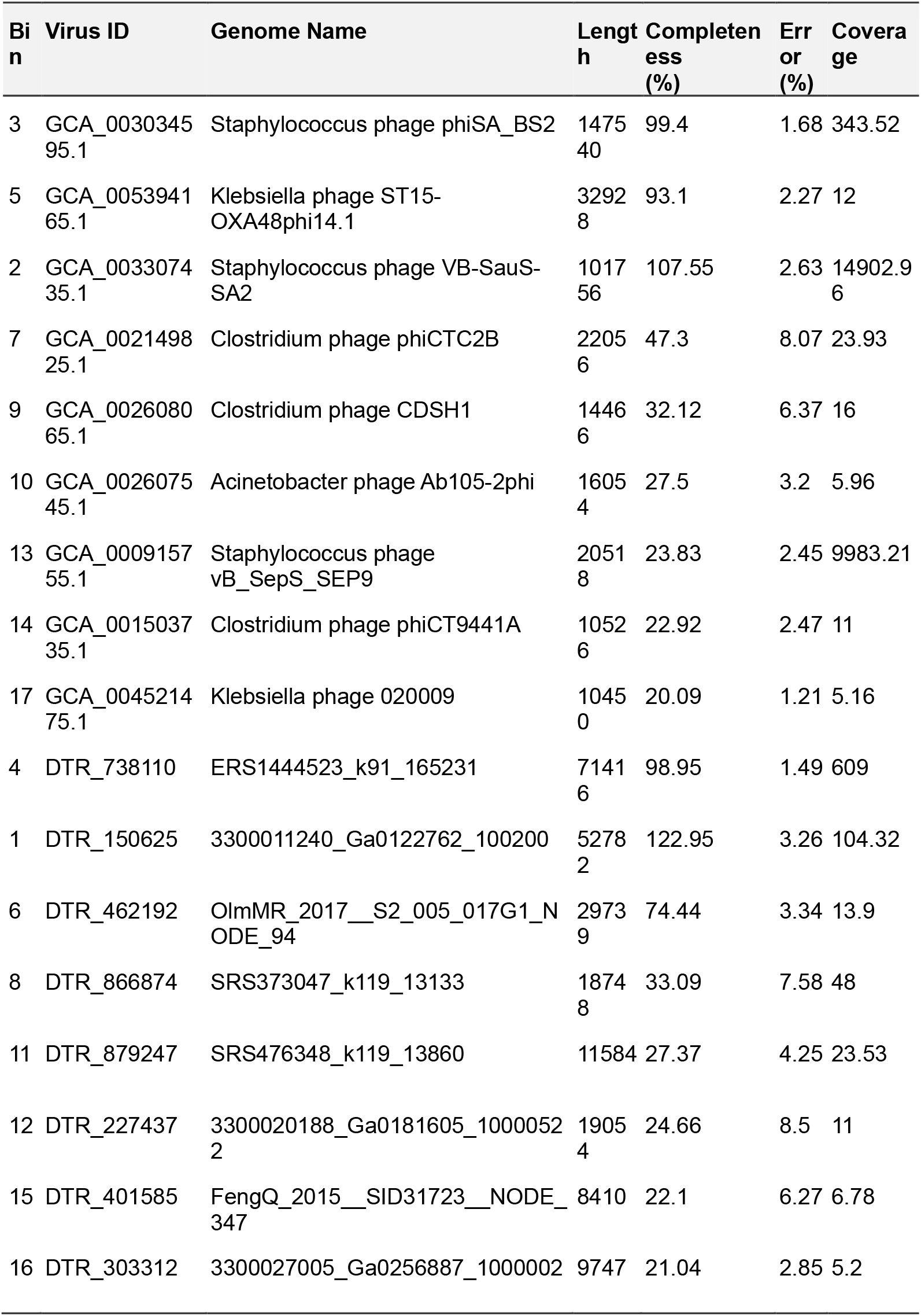
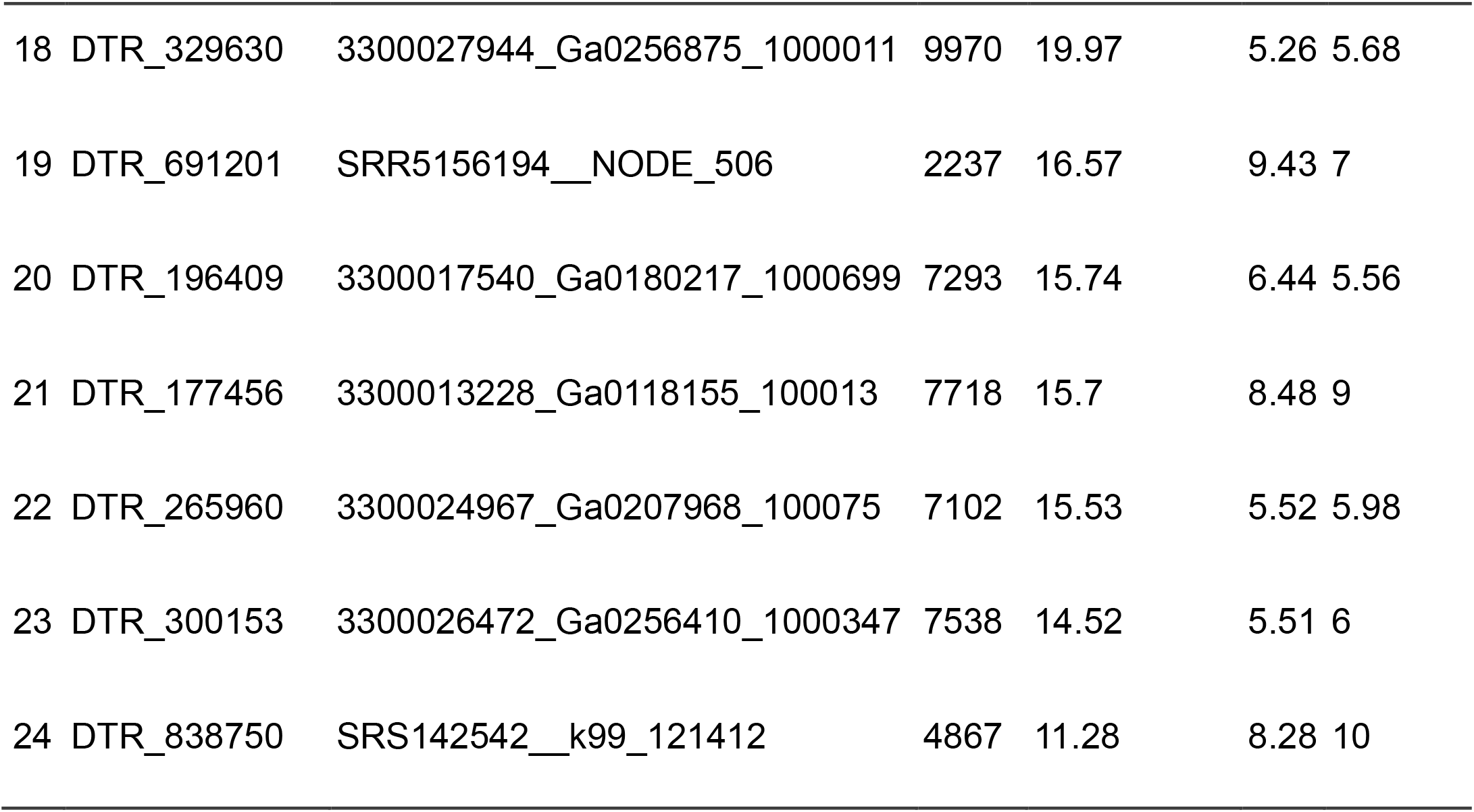
Binning results for sample RSMJ51Sa. Parameters including possible relative strain hit, bin size (length), completeness and other features are presented in corresponding columns.

| Bin | Virus ID | Genome Name | Length | Completeness (%) | Error (%) | Coverage |
| --- | --- | --- | --- | --- | --- | --- |
| 3 | GCA_003034595.1 | Staphylococcus phage phiSA_BS2 | 147540 | 99.4 | 1.68 | 343.52 |
| 5 | GCA_005394165.1 | Klebsiella phage ST15-OXA48phi14.1 | 32928 | 93.1 | 2.27 | 12 |
| 2 | GCA_003307435.1 | Staphylococcus phage VB-SauS-SA2 | 101756 | 107.55 | 2.63 | 14902.96 |
| 7 | GCA_002149825.1 | Clostridium phage phiCTC2B | 22056 | 47.3 | 8.07 | 23.93 |
| 9 | GCA_002608065.1 | Clostridium phage CDSH1 | 14466 | 32.12 | 6.37 | 16 |
| 10 | GCA_002607545.1 | Acinetobacter phage Ab105-2phi | 16054 | 27.5 | 3.2 | 5.96 |
| 13 | GCA_000915755.1 | Staphylococcus phage vB_SepS_SEP9 | 20518 | 23.83 | 2.45 | 9983.21 |
| 14 | GCA_001503735.1 | Clostridium phage phiCT9441A | 10526 | 22.92 | 2.47 | 11 |
| 17 | GCA_004521475.1 | Klebsiella phage 020009 | 10450 | 20.09 | 1.21 | 5.16 |
| 4 | DTR_738110 | ERS1444523_k91_165231 | 71416 | 98.95 | 1.49 | 609 |
| 1 | DTR_150625 | 3300011240_Ga0122762_100200 | 52782 | 122.95 | 3.26 | 104.32 |
| 6 | DTR_462192 | OlmMR_2017__S2_005_017G1_NODE_94 | 29739 | 74.44 | 3.34 | 13.9 |
| 8 | DTR_866874 | SRS373047_k119_13133 | 18748 | 33.09 | 7.58 | 48 |
| 11 | DTR_879247 | SRS476348_k119_13860 | 11584 | 27.37 | 4.25 | 23.53 |
| 12 | DTR_227437 | 3300020188_Ga0181605_10000522 | 19054 | 24.66 | 8.5 | 11 |
| 15 | DTR_401585 | FengQ_2015__SID31723__NODE_347 | 8410 | 22.1 | 6.27 | 6.78 |
| 16 | DTR_303312 | 3300027005_Ga0256887_1000002 | 9747 | 21.04 | 2.85 | 5.2 |
| 18 | DTR_329630 | 3300027944_Ga0256875_1000011 | 9970 | 19.97 | 5.26 | 5.68 |
| 19 | DTR_691201 | SRR5156194__NODE_506 | 2237 | 16.57 | 9.43 | 7 |
| 20 | DTR_196409 | 3300017540_Ga0180217_1000699 | 7293 | 15.74 | 6.44 | 5.56 |
| 21 | DTR_177456 | 3300013228_Ga0118155_100013 | 7718 | 15.7 | 8.48 | 9 |
| 22 | DTR_265960 | 3300024967_Ga0207968_100075 | 7102 | 15.53 | 5.52 | 5.98 |
| 23 | DTR_300153 | 3300026472_Ga0256410_1000347 | 7538 | 14.52 | 5.51 | 6 |
| 24 | DTR_838750 | SRS142542__k99_121412 | 4867 | 11.28 | 8.28 | 10 |

Table 2 represents the binning results for sample RSMJ51Sa. High quality MAGs include bin 3 (*Staphylococcus* phage phiSA_BS2; accession GCA_003034595.1),5 (*Klebsiella* phage ST15-OXA48phi14; accession GCA_005394165.1), 2 (*Staphylococcus* phage VB-SauS-SA2; GCA_003307435.1), 4 and 1 (undetected NCBI matches).

Fig.2 represents the phylogrouping of high-quality MAGs against selected reference phage genomes retrieved from GenBank. The datasets show that various phage species were co-isolated from the enrichment and filtration process, which were then recovered from the overnight cultivation of the *Staphylococcus* host. The diversity includes phages targeting *Staphylococcus, Escherichia, Acinetobacter, Klebsiella* and *Vibrio* spp., among others.

**Fig. 2.**
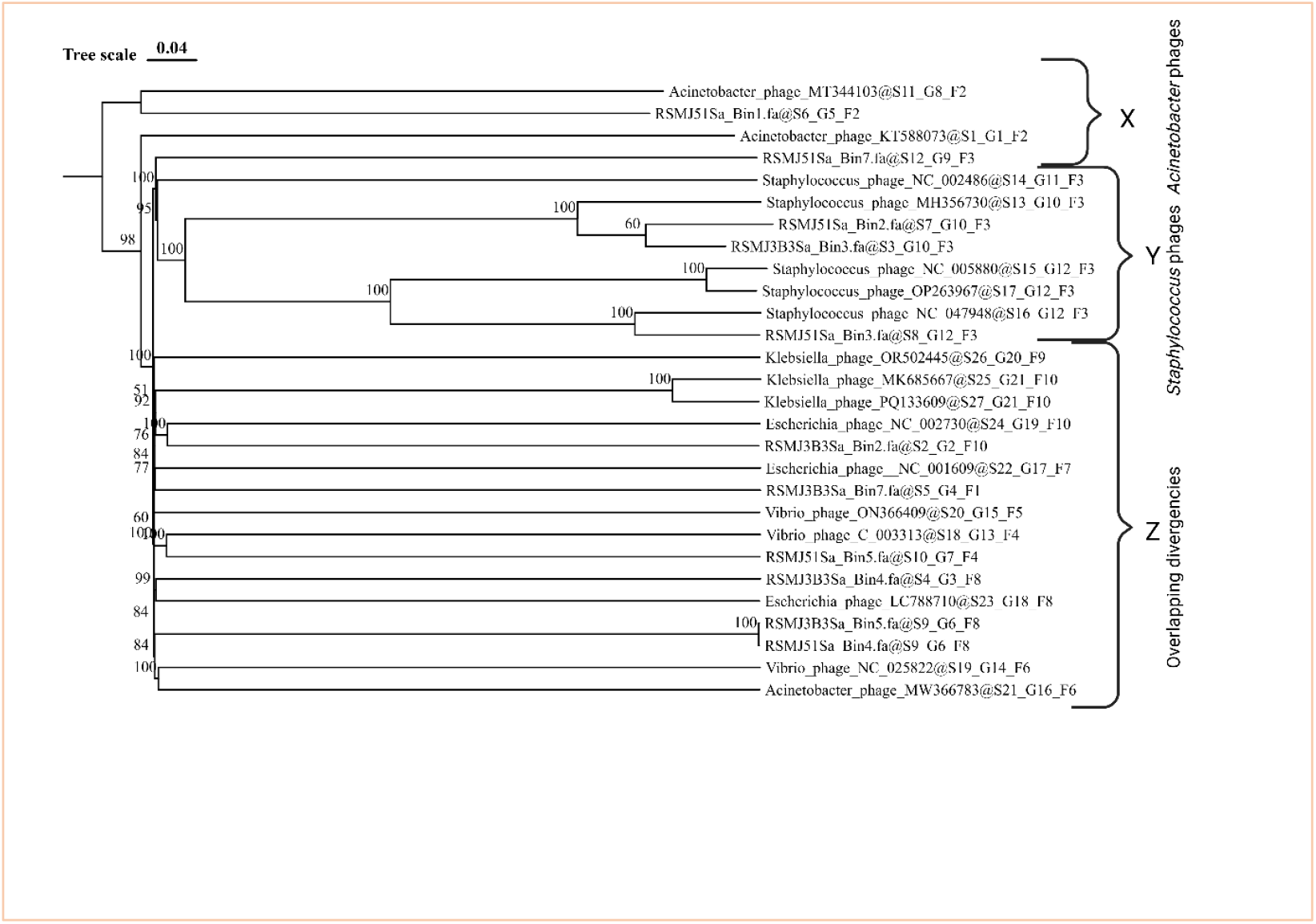
The phylogenomic GBDP tree was constructed using the D4 FASTME formula, achieving an average support value of 90. The numbers displayed above the branches represent GBDP pseudo-bootstrap support values derived from 100 replications. The branch lengths in the resulting VICTOR trees are proportionally scaled according to the specific distance formula applied.

### PhageScope annotation datasets

Datasets from PhageScope annotation of high-quality anti-ESKAPE phages are presented in Fig. 3 (a, b. c, d). The dataset in this figure includes selected high-quality MAGs with relevance to ESKAPE and/or anti-CRISPR/Cas potential. The datasets emphasize the genomic differences across the MAGs in which *epidermidis Staphylococcus epidermidis* phage MAG (RSMJ3B3Sa bin 3) possesses many terminator genes, suggestive of its host gene expression regulation effects. The second MAG (RSMJ51Sa bin 1) is an *Acinetobacter* phage with significant lysis machinery comprising of holin and lysozyme (endolysin), the most effective facets required to damage bacterial membranes and suppress their multiplication [6]. The *Klebsiella* phage MAG (RSMJ51Sa bin 5) carries only two transcription terminator genes but by far more transmembrane proteins, suggestively promoting effective host membrane attachment and disruption [7]. Information with additional PhageScope-annotated high-quality MAG datasets is presented in Table 3. These datasets show that the first sample RSMJ3B3Sa MAG, bin 3 belongs to *Staphylococcus* phage, annotated as a temperate phage of *S. epidermidis*. Bin 4 matched *Vibrio* phage and was recognized as a virulent pathogen of *Vibrio alginolyticus*.

**Table 3.**
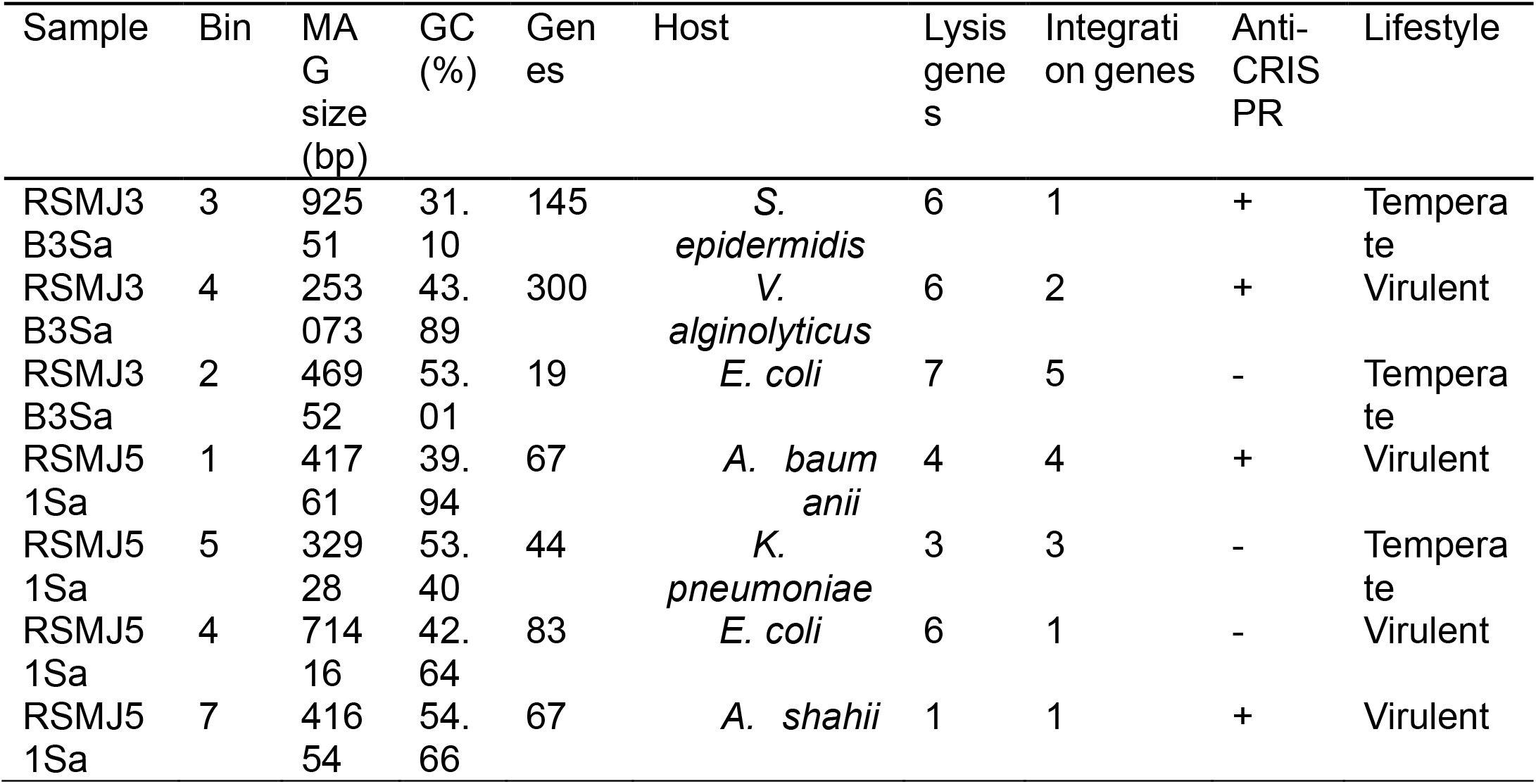
A summary of PhageScope-annotated MAGs. All the MAGs were taxonomically assigned to Caudoviricetes, a class of tailed bacteriophages. Features relating to anti-CRISPR are indicated as “+”, meaning that the genome harbors anti-CRISPR genes, or “-“to imply that the phage genome carries no anti-CRISPR genes. Virulent lifestyle implies that the phage is obligately lytic while temperate implies that the phage could alternate between lytic and lysogenic forms.

**Fig. 3.**
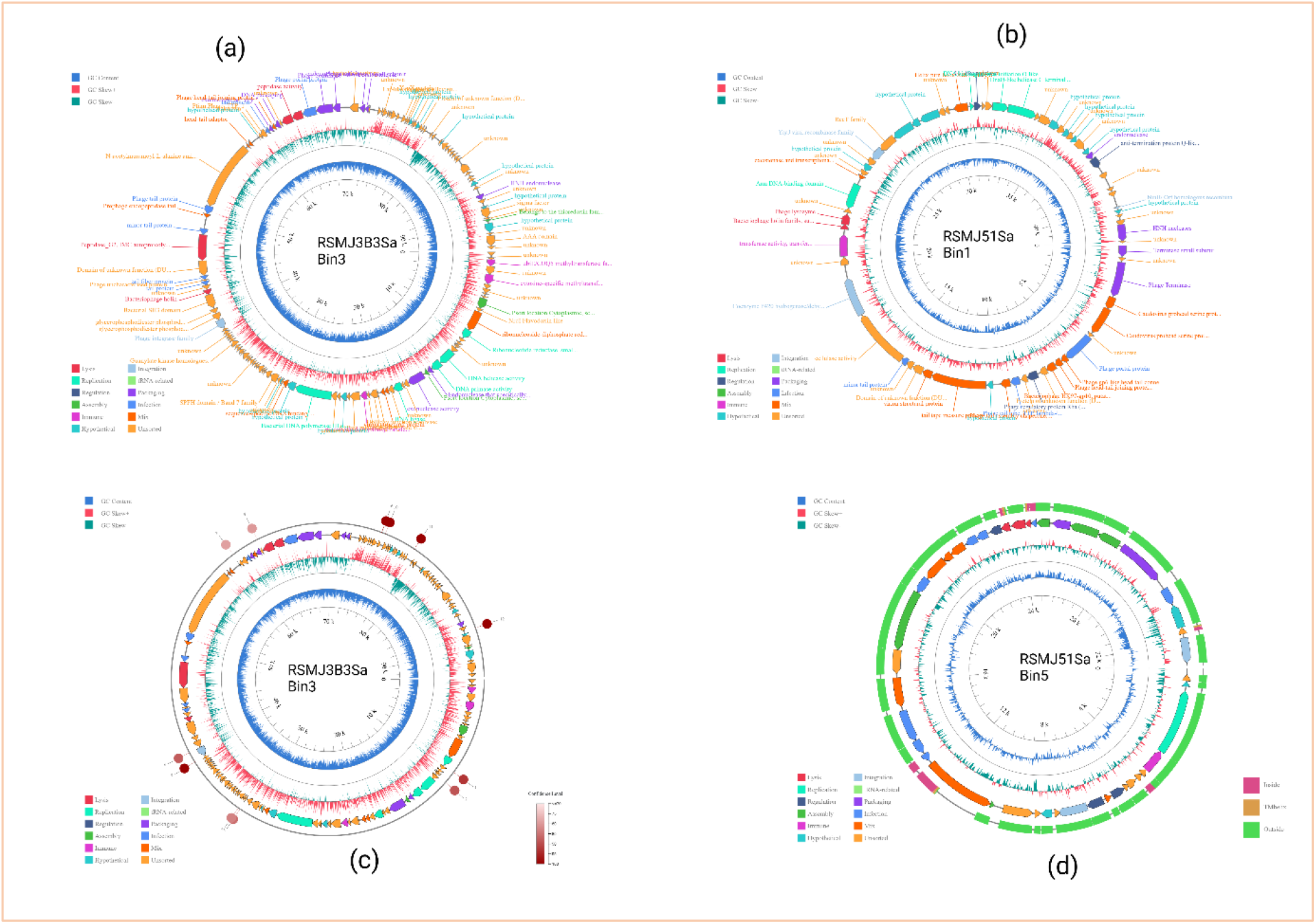
*Staphylococcus Acinetobacter*, and *Klebsiella* phage genomes. The circular genome diagrams illustrate anti-ESKAPE phages with varying functional strengths. (a) Complete genome annotation of RSMJ3B3Sa bin 3, a *Staphylococcus* phage. (b) Full genome annotation of RSMJ51Sa bin 1, an *Acinetobacter* phage. (c) The *Staphylococcus* phage RSMJ3BSa bin 3 highlights significant transcription terminator proteins. (d) The *Klebsiella* phage RSMJ51Sa bin 5 features numerous open reading frames (ORFs) associated with transmembrane proteins.

On the other hand, MAG databases of the second sample RSMJ51Sa include bin 1, which was annotated as *Acinetobacter* phage and as a virulent pathogen of *A. baumanii*, with possession of an anti-CRISPR gene positioned from 41,662-41,760 bp of the genome. The second phage anti-ESKAPE MAG corresponded to bin 5, which was annotated as temperate. Details of MAG features from PhageScope are presented in Table 3.

## ETHICS STATEMENT

This work did not involve animal or human subjects. However, it was approved by the Kampala International University (KIU) Research Ethical Committee (REC) through the approval number KIU-2024-435.

## AUTHORSHIP ROLES AND RESPONSIBILITIES

**Jackim Nabona:** Conceptualization, Methodology, Software. Writing, Original draft preparation, **Samweli Bahati**: Software, Data curation, Writing, Original draft preparation. **Abdalah Makaranga**: Methodology, Visualization, Investigation. **Chinyere Nkemjika Anyanwu:** Writing-Reviewing and Editing Methodology, Supervision. **R Neel:** Writing-Reviewing and Editing. **Emmanuel Eilu**: Supervision, Writing-Reviewing and Editing, **Deogratius Mark**: Conceptualization, Methodology, Writing-Reviewing and Editing. **Reuben S. Maghembe:** Conceptualization, Methodology, Software. Writing, Original draft preparation, Supervision

## ACKNOWLEDGEMENTS

This research did not receive any specific grant from funding agencies in the public, commercial, or not-for-profit sectors

## REFERENCES

[1] C.E. Aruwa, T. Chellan, N.W. S’thebe, Y. Dweba, S. Sabiu, ESKAPE pathogens and associated quorum sensing systems: New targets for novel antimicrobials development, Health Sciences Review 11 (2024) 100155. 10.1016/j.hsr.2024.100155.

[2] L. Daruka, M.S. Czikkely, P. Szili, Z. Farkas, D. Balogh, G. Grézal, E. Maharramov, T.-H. Vu, L. Sipos, S. Juhász, A. Dunai, A. Daraba, M. Számel, T. Sári, T. Stirling, B.M. Vásárhelyi, E. Ari, C. Christodoulou, M. Manczinger, M.Z. Enyedi, G. Jaksa, K. Kovács, S. van Houte, E. Pursey, L. Pintér, L. Haracska, B. Kintses, B. Papp, C. Pál, ESKAPE pathogens rapidly develop resistance against antibiotics in development in vitro, Nature Microbiology (2025). 10.1038/s41564-024-01891-8.

[3] F. Newberry, P. Shibu, T. Smith-Zaitlik, M. Eladawy, A.L. McCartney, L. Hoyles, D. Negus, Lytic bacteriophage vB_KmiS-Kmi2C disrupts biofilms formed by members of the Klebsiella oxytoca complex, and represents a novel virus family and genus, Journal of Applied Microbiology 134 (2023) xad079. 10.1093/jambio/lxad079.

[4] E.M. Townsend, L. Kelly, L. Gannon, G. Muscatt, R. Dunstan, S. Michniewski, H. Sapkota, S.J. Kiljunen, A. Kolsi, M. Skurnik, T. Lithgow, A.D. Millard, E. Jameson, Isolation and Characterization of Klebsiella Phages for Phage Therapy, PHAGE 2 (2021) 26–42. 10.1089/phage.2020.0046.

[5] P. Hyman, Phages for Phage Therapy: Isolation, Characterization, and Host Range Breadth, Pharmaceuticals 12 (2019). 10.3390/ph12010035.

[6] M. Khan Mirzaei, Md.A.A. Khan, P. Ghosh, Z.E. Taranu, M. Taguer, J. Ru, R. Chowdhury, Md.M. Kabir, L. Deng, D. Mondal, C.F. Maurice, Bacteriophages Isolated from Stunted Children Can Regulate Gut Bacterial Communities in an Age-Specific Manner, Cell Host & Microbe 27 (2020) 199-212.e5. 10.1016/j.chom.2020.01.004.

[7] G.S. Abeysekera, M.J. Love, S.H. Manners, C. Billington, R.C.J. Dobson, Bacteriophage-encoded lethal membrane disruptors: Advances in understanding and potential applications, Frontiers in Microbiology 13 (2022). 10.3389/fmicb.2022.1044143.

[8] R.N. Beaudoin, D.R. DeCesaro, D.L. Durkee, S.E. Barbaro, ISOLATION OF A BACTERIOPHAGE FROM SEWAGE SLUDGE AND CHARACTERIZATION OF ITS BACTERIAL HOST CELL, in: 2007. https://api.semanticscholar.org/CorpusID:37901910.

[9] R.S. Maghembe, F.P. Mdoe, A. Makaranga, J.A. Mpemba, D. Mark, C. Mlay, E.A. Moto, A.G. Mtewa, Complete genome sequence data of Priestia megaterium strain MARUCO02 isolated from marine mangrove-inhabited sediments of the Indian Ocean in the Bagamoyo Coast, Data in Brief 48 (2023) 109119. 10.1016/j.dib.2023.109119.

[10] J. Lu, F.P. Breitwieser, P. Thielen, S.L. Salzberg, Bracken: estimating species abundance in metagenomics data, PeerJ Computer Science 3 (2017) e104. 10.7717/peerj-cs.104.

[11] D. Li, C.-M. Liu, R. Luo, K. Sadakane, T.-W. Lam, MEGAHIT: an ultra-fast single-node solution for large and complex metagenomics assembly via succinct de Bruijn graph, Bioinformatics 31 (2015) 1674–1676. 10.1093/bioinformatics/btv033.

[12] A. Chklovski, D.H. Parks, B.J. Woodcroft, G.W. Tyson, CheckM2: a rapid, scalable and accurate tool for assessing microbial genome quality using machine learning, Nature Methods 20 (2023) 1203–1212. 10.1038/s41592-023-01940-w.

[13] J.P. Meier-Kolthoff, M. Göker, VICTOR: genome-based phylogeny and classification of prokaryotic viruses, Bioinformatics 33 (2017) 3396–3404. 10.1093/bioinformatics/btx440.

[14] J.P. Meier-Kolthoff, A.F. Auch, H.-P. Klenk, M. Göker, Genome sequence-based species delimitation with confidence intervals and improved distance functions, BMC Bioinformatics 14 (2013) 60. 10.1186/1471-2105-14-60.

[15] V. Lefort, R. Desper, O. Gascuel, FastME 2.0: A Comprehensive, Accurate, and Fast Distance-Based Phylogeny Inference Program, Molecular Biology and Evolution 32 (2015) 2798–2800. 10.1093/molbev/msv150.

[16] J.S. Farris, Estimating Phylogenetic Trees from Distance Matrices, The American Naturalist 106 (1972) 645–668.

[17] G. Yu, Using ggtree to Visualize Data on Tree-Like Structures, Current Protocols in Bioinformatics 69 (2020) e96. 10.1002/cpbi.96.

[18] R.H. Wang, S. Yang, Z. Liu, Y. Zhang, X. Wang, Z. Xu, J. Wang, S.C. Li, PhageScope: a well-annotated bacteriophage database with automatic analyses and visualizations, Nucleic Acids Research 52 (2024) D756–D761. 10.1093/nar/gkad979.

